## Supplemental Info for "Low-cost drug discovery with engineered *E. coli* reveals an anti-mycobacterial activity of benazepril"

### Included in this file:

| | |
|---|---|
| **ALR** | Alanine Racemase |
| **DCS** | D-Cycloserine |
| ***Mtb*** | *Mycobacterium tuberculosis* |
| **TESEC** | Target-Essential Surrogate *E. coli* |

**Table S1 | Abbreviations used in this text.**

### SUPPLEMENTARY RESULTS

#### Construction of an efflux-deficient TESEC host with minimal growth perturbation

The TolC outer membrane channel is required for the function of many efflux systems that contribute to antibiotic resistance in *E. coli*(1). While not all efflux activity is TolC-dependent, most clinically relevant forms of resistance are attributed to TolC and the many inner membrane permeases with which it interacts(2).

Deletion mutants of *tolC* suffer from reduced growth rates and abnormal cell morphology. Vega and Young showed that these defects were caused by an accumulation of enterobactin in the periplasm and could be minimized by a compensating deletion in the isochorismate synthase *entC*(3).

To increase drug sensitivity while minimizing other phenotypic perturbations, we introduced the *tolC* and *entC* deletions in the TESEC Host (Fig S1). Deletion of *tolC* was associated with an extended lag phase but no reduction in growth rate or saturation density (Fig. S1A). Cell morphology of the Δ*tolC* strain was elongated and aberrant as evaluated by phase microscopy (Fig. S1B). Introduction of the compensating *entC* deletion in restored a wild-type rod shape and reduced, but did not eliminate, the extended lag phase.

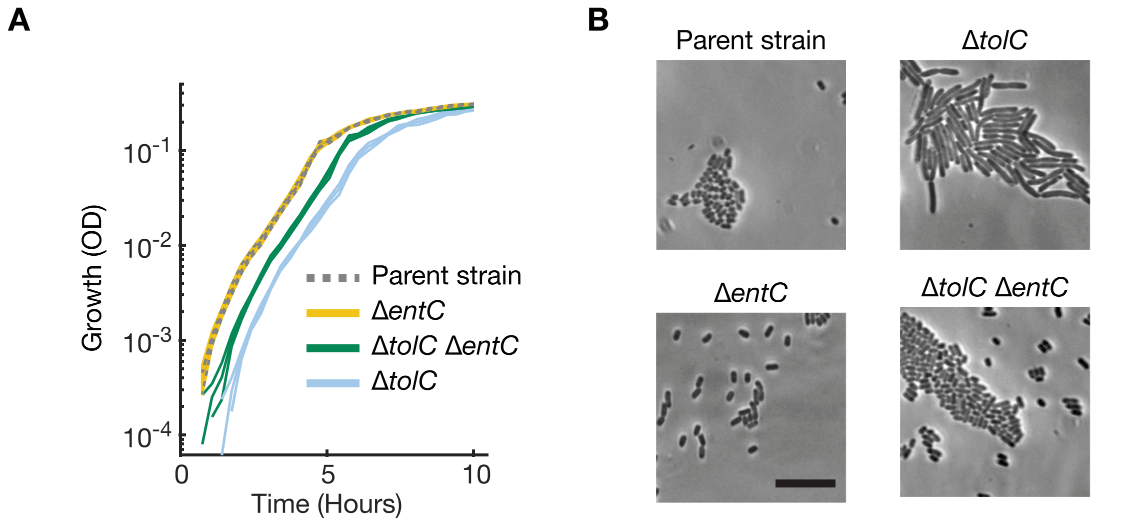

**Supplementary Figure S1 | Deletion of *entC* partially restores growth and morphology defects of the efflux-deficient *tolC* deletion mutant.** A) The indicated deletions were introduced to the TESEC parent strain BW25113(4). Overnight cultures of cells were diluted to 0.001 in defined medium and dispensed at 180 µL into 96-well, flat-bottom microplates. Cells were grown at 37 °C with shaking in a TECAN Infinite plate reader. Individual growth traces are shown for 3 biological replicates. B) Phase-contrast microscopy images (63x magnification) of strains with the indicated deletions. Cells were grown to saturation in defined medium. Scale bar is 10 µm.

#### Evaluation of anti-mycobacterial activity of benazepril by spot plating

We sought to characterize the activity of benazepril as a bactericidal in addition to it's effect as a bacteriostatic described in the main text. Standard serial dilutions and spot plating *of M. smegmatis* $mc^2155$ showed significant killing caused by both DCS and benazepril in the millimolar range (Fig S2).

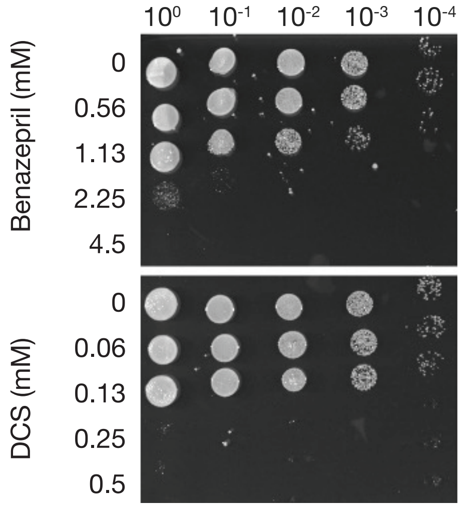

**Supplementary Figure S2 | Benazepril and DCS show bactericidal activity near their growth-inhibitory concentrations.** *M. smegmatis* mc²155 cultures in Middlebrook 7H9 medium (Sigma M0178) were exposed to DCS (0-0.5 mM) or benazepril (0-4.5 mM) for 48 hours. Serial dilutions were prepared of the bacterial cultures and spotted as 1 µL aliquots onto Middlebrook 7H9 agar plates. A photo was taken after 48 hours of incubation at 37 °C. The image is representative of three independent experiments.

**Validation of an in vitro coupled assay for inhibition of ALR with benazepril**

Our in vitro assay for ALR activity relies on a coupled reaction to alanine dehydrogenase(5). Metabolic inhibitors, particularly substrate analogs, may act on multiple targets within a metabolic pathway if pathway intermediates are structurally similar. DCS, for example, acts both on ALR and the downstream D-alanine:D-alanine ligase(6).

We therefore sought to confirm that benazepril does not inhibit L-alanine dehydrogenase directly. Neither DCS nor benazepril was active against L-alanine dehydrogenase under conditions similar to the full ALR assay.

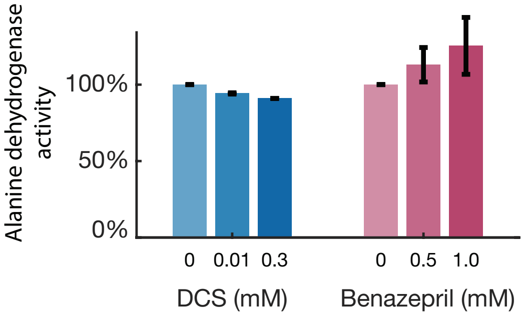

**Supplementary Figure S3 | ALR inhibitors do not affect the second enzyme of the in vitro coupled ALR assay.** L-alanine dehydrogenase from *Bacillus subtilis* (Merck A7653) was prepared at 50 U/µL in reaction buffer matching the ALR inhibition assays (20 mM Tris, 100 mM NaCl, pH 8.0, 20 mM NAD+). Addition of 5 mM L-alanine and benazepril or DCS at the indicated concentrations did not significantly reduce the rate of NADH production, as measured by time-course fluorescence measurements. Error bars represent the standard deviations of 3 biological replicates.
